# Field application of the geometric framework reveals a multistep strategy of nutrient regulation in a leaf-miner

**DOI:** 10.1101/777367

**Authors:** Mélanie J.A. Body, Spencer T. Behmer, Pierre-François Pelisson, Jérôme Casas, David Giron

**Author notes:** Author for correspondence: Dr. David Giron, Address: Institut de Recherche sur la Biologie de l’Insecte, UMR 7261, CNRS/Université François-Rabelais, Parc Grandmont, 37200 Tours, France. Author Contributions: MJAB and DG conceived and designed the experiments. MJAB, PFP and DG performed the experiments. MJAB, STB, JC and DG analyzed and interpreted the data. MJAB, STB and DG wrote the manuscript; all authors provided editorial advice.

## Abstract

Animals have evolved a vast array of behavioral and physiological strategies that allow them to achieve a nutritionally balanced diet. Plants as food for herbivores are often considered suboptimal, but phytophagous insects can employ pre- and post-ingestive mechanisms and/or symbiotic associations to help overcome food nutritional imbalances. This is particularly crucial for permanent multivoltine leaf-miner insects such as the caterpillar *Phyllonorycter blancardella* which completes development within a restricted area of a single leaf and use deciduous leaves to fuel growth and reproduction even under senescing autumnal conditions. Using the geometric framework for nutrition under natural field conditions, we show that this insect has multiple strategies to deal with inadequate food supply from the plant. First, larvae manipulate the protein-sugar content of both normal, photosynthetically active, and senescing, photosynthetically inactive, leaf tissues. Control of nutritional homeostasis of mined tissues is however higher for late instars, which differ from younger larval instars in their feeding mode (fluid-*vs.* tissue-feeder). Second, slight differences in the protein-sugar environment remain between mined tissues on green and yellow leaves despite this manipulation of the leaf physiology. This insect uses post-ingestive mechanisms to achieve similar body protein, sugar and lipid composition. This study demonstrates, for the first time under natural conditions, the ability of an insect herbivore to practice a combination of pre- and post-ingestive compensatory mechanisms to attain similar growth and metabolic outcomes in fundamentally different nutritional environments. Additionally, a comparison of larval nutritional requirements of 117 species from various insect groups further reinforces the hypothesis of a close association between *P. blancardella* and endosymbiotic bacteria for nutritional purposes.

## Introduction

Feeding on plant tissues is challenging for vertebrate and insect herbivores as this food source is often considered nutritionally suboptimal due to their nitrogen-limited base (Mattson 1980; White 1993; Schoonhoven et al. 2005). This nitrogen limitation has been recorded in herbivores ranging from large mammals (e.g. impalas, springboks, blesboks, spider monkeys) to insects (e.g. grasshoppers, locusts, caterpillars, leaf-miners) (Simpson et al. 2002; Van Zyl and Ferreira 2003; Felton et al. 2009; Joern et al. 2012; Barbehenn et al. 2013; Roeder and Behmer 2014; Body et al. in prep; see e.g. Rothman et al. 2011 for exception on gorillas). Additionally, the nutritional environment is frequently highly variable both in space and time (Joern et al. 2012). Likewise, nutritional requirements of animals are multidimensional and change qualitatively and quantitatively as an individual grows, develops, becomes reproductively active, then senesces (Schoonhoven et al. 2005; Rothman et al. 2008, 2011; Behmer 2009; Raubenheimer et al. 2009). Thus, food intake for any given life-stage is not necessarily matched to life-history trait requirements for that life-stage. However, herbivores can employ a suite of pre- and post-ingestive mechanisms to address this nutritional mismatch (Simpson and Raubenheimer 1993; Behmer 2009; Raubenheimer et al. 2009).

Different strategies can allow herbivores to regulate their nutrient intake pre-ingestion (Figure 1). (*i*) Herbivores can select a food source that matches its nutritional requirement. Although this case is ideal, it is most likely to be rare in nature. (*ii*) They can also feed on an unbalanced food source and compromise by overeating one nutrient while undereating another nutrient (Figure 1b) (Rothman et al. 2011). (*iii*) An herbivore can reach its intake target by regulating the amount of an individual plant part that is eaten, feeding from a range of different plants or, more likely, through a combination of these two mechanisms (Behmer 2009; Felton et al. 2009; Rothman et al. 2011). In the case of a specialist herbivore, the same combination of mechanisms can be used, but mixing occurs by feeding on vegetative tissues of different age classes (e.g., young *vs*. old) or switching between individual plants within the same species or family (Figure 1c) (Behmer 2009). Switching between different food sources can occur at any timescale, ranging from bites to days, and the rate at which switching occurs is determined by the costs associated with such behaviors.

**Figure 1.**
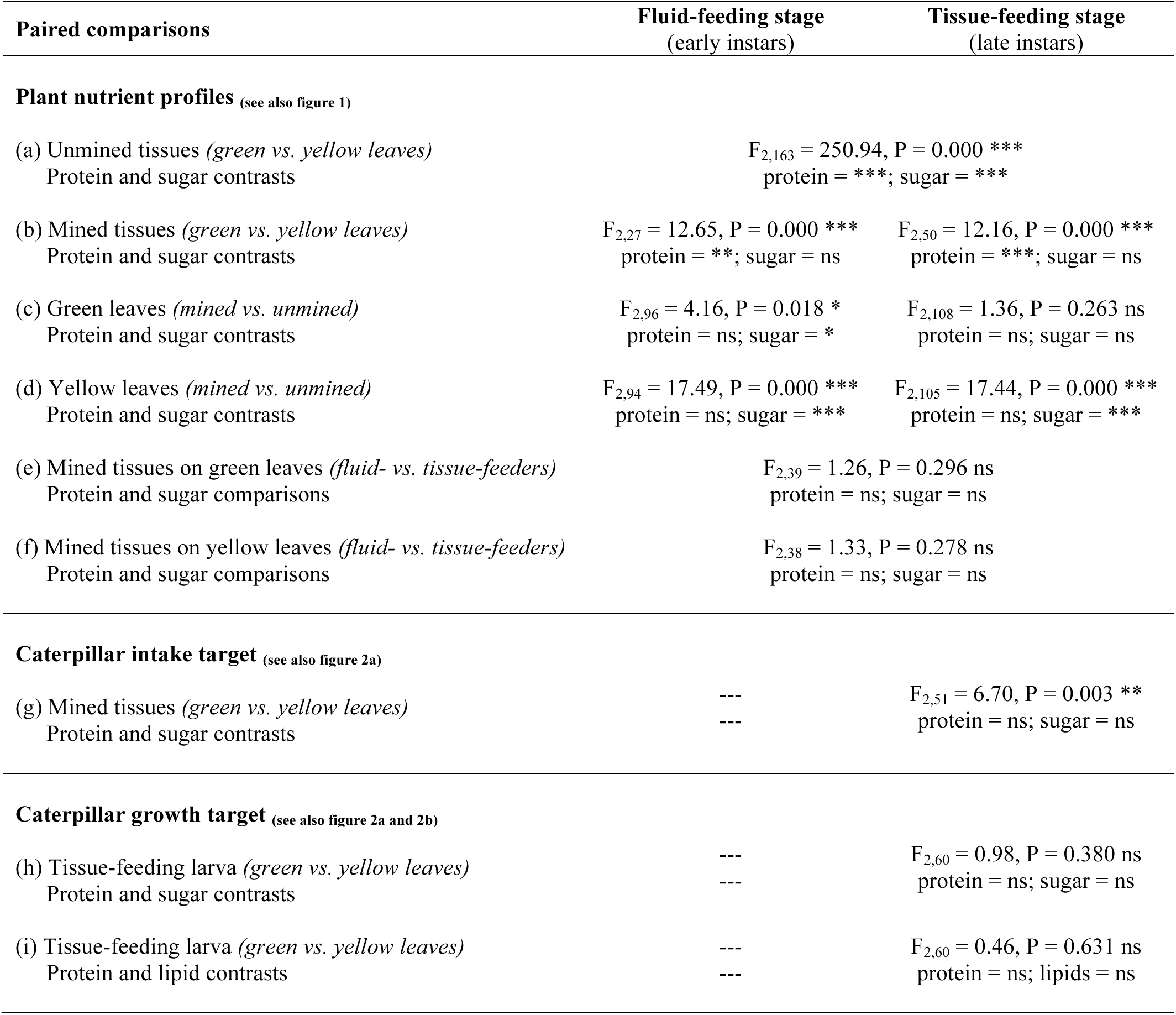
Geometric framework (GF). Panel (a) shows an herbivorous insect that has a protein-carbohydrate (P:C) intake target ratio of 1:1 (black target). The insect can reach this intake target by consuming an optimal balanced food (dark green line; ideal scenario). Panel (b) depicts a situation in which there is access only to a single unbalanced food that contains protein and carbohydrate in a 1:2 ratio (green line). The insect herbivore is unable to reach its intake target (black target) leading to nutritional compromises characterized by three options: (*i*) feed until it meets its requirement for carbohydrate (light green circle) but suffers a deficit in protein (the amount of the deficit is the length of the light green solid line); (*ii*) feed until it meets its requirement for protein (dark green circle) but suffers an excess of carbohydrate (the amount of the excess is the length of the dark green solid line); or (*iii*) feed to some intermediate point (white circle) so that the experienced excesses (represented by the dark green dotted line) and deficits (represented by the light green dotted line) are less extreme. The extent to which an insect herbivore (or any animal) overeats one nutrient while undereats another represents a compromise employed by insect. Panel (c) shows an insect herbivore that can reach his intake target (black target) by switching adequately between nutritionally unbalanced, suboptimal but complementary foods without accumulating extreme nutrient excesses or deficits. In this example, there are two foods, one with a P:C ratio of 2:1 (light green line) and the other with a P:C ratio of 1:2 (dark green line). Each dotted line represents an individual meal, with length correlated with meal size. The shaded area represents the available nutrient space, as defined by the nutrient rails of the two foods. Panel (d) depicts a hypothetic situation where an endophagous insect is able to manipulate its food source to achieve the same nutrient intake (black target) when the nutrient quantity and/or quality changes over the course of its development or lifecycle. For example, when the leaf-miner starts its development on photosynthetically active green leaves (green line) but completes it after senescence occurred (senescing photosynthetically inactive leaves; yellow line). The shaded area represents the available nutrient space, as defined by the nutrient rails of the two extreme foods. Adapted from Raubenheimer and Simpson 1999; Behmer 2009.

In the context of insect nutritional ecology, endophytophagous organisms such as gall-inducing and permanent leaf-mining insects are peculiar because they simultaneously live in, and eat their food; they also live in a restricted nutritional environment (Stone and Schönrogge 2003; Giron et al. 2016). As a consequence, while endophagous insects by their feeding habit secure their nutrition and shelter for either shorter or longer periods of their life history, they are also trapped in a very restricted area within plant tissues with no possibilities to switch between plants or leaves if their food source varies in quantity and/or quality. Endophagous insects have evolved specific feeding strategies to deal with this challenging environment by altering the plant morphology and physiology for their own benefits (Stone and Schönrogge 2003; Giron et al. 2016). This allows them to manipulate their host plant in a way to best meet their needs, including counteracting plant defenses, and compensating for variation in food nutritional composition (Figure 1d) (Stone and Schönrogge 2003).

Gall-inducing insects have long been known to alter the plant morphology and physiology for their own benefits, but data on the nutritional needs of leaf-miners, and their potential capacity to modify the plant to meet nutritional needs, have remained however scarce (Giron et al. 2016). Recently, the spotted tentiform leaf-miner *Phyllonorycter blancardella* has been shown to manipulate plant vegetative tissue in a particularly spectacular way (Giron et al. 2007; Body et al. 2013; Zhang et al. 2016; Body et al. in prep). This leaf-miner spends its entire larval life-cycle within a small area of a single leaf with no possibilities to extend its nutritional micro-environment, or to switch between plants or leaves in case of inadequate food supply (Body et al. 2015). As a multivoltine species, different generations of this insect experience, over the course of a season, leaves with different nutritional profiles (Body et al. 2013; Body et al. in prep). Within a single generation, insects also interact with leaves offering different ratios of nutrients. For instance, the last generation has to face adverse autumnal conditions where senescing yellow leaves represent for the developing larva a poor and declining source of nutrients with a lower sugar content relative to green leaves (Body et al. 2013; Body et al. in prep). However, to face these constraints, *P. blancardella* and several other leaf-miner species can prevent mined tissues from senescing (inducing a ‘green-island’ phenotype) through a manipulation of the plant cytokinin profile (Engelbrecht et al. 1969; Giron et al. 2007; Body et al. 2013; Gutzwiller et al. 2015; Zhang et al. 2016). Recently, a bacterial symbiont (most likely *Wolbachia*) mediated cytokinin release to the plant has been revealed in this system, both in senescing and photosynthetically active leaf tissues (Giron et al. 2007; Kaiser et al. 2010; Body et al. 2013; Gutzwiller et al. 2015; Zhang et al. 2017). The correlation between the green-island phenotype and *Wolbachia* infections has also been highlighted in numerous species of Gracillariidae leaf-mining moths (Gutzwiller et al. 2015). Recent studies on this system also showed a strong reprogramming of the plant phytohormonal balance (cytokinins, jasmonic acid, salicylic acid, abscisic acid) associated with a mitigation of plant direct and indirect defense, inhibition of leaf senescence and increased nutrient mobilization (sugars and amino acids) (Body et al. 2013; Zhang et al. 2016; Body et al. in prep). Cytokinins are known to regulate plant sugar metabolism, allowing insects to control for the sugar profile of mined tissues (Body et al. 2013; Body et al. in prep). However, their impact on other key insect fitness related nutrients such as plant proteins is poorly documented. Additionally, *P. blancardella* exhibits two distinct feeding modes. First instars are fluid-feeders, while last instars are tissue-feeders that selectively eat mesophyll cells (Body et al. 2015). Previous studies on the impact of fluid- and tissue-feeding larvae on the individual sugar and (protein-bound and free) amino acid contents showed a differential ability to manipulate their nutritional environment, the control being finely-tuned only by tissue-feeding instars (Body 2013; Body et al. in prep). However, the extent to which the nutrient modifications in mined tissues is beneficial for the insect requires quantifying soluble sugar and protein consumption in caterpillars, and documenting how these nutrients are utilized for growth (e.g., levels of protein, sugar and lipid in the body) (Simpson and Raubenheimer 1993; Behmer 2009; Raubenheimer et al. 2009). Understanding this requires investigating the ability of both feeding modes to manipulate the protein and soluble sugar profiles of the host-plant *Malus domestica*, including comparisons of photosynthetically active and senescing plant tissues.

Such questions require a nutritional approach initially developed under highly controlled lab conditions to investigate insect nutritional ecology to link food nutrient content, nutrient consumption, and nutrient allocation to growth. This approach deemed the ‘Geometric Framework’ for nutrition (henceforth GF), provides a theoretical unifying framework leading to a deep understanding of behavioral and physiological mechanisms underlying nutritional homeostasis and how the nutritional environment of animals impact their performances (Simpson and Raubenheimer 1993; Raubenheimer and Simpson 1999; Behmer 2009; Raubenheimer et al. 2009). First, the GF depicts an animal as living in a multidimensional ‘nutrient space’, where foods can be defined by the amounts and ratios of their nutritional constituents (typically macronutrients). Second, if the nutritional value of a given food can be measured, and if food consumption can be quantified, the GF makes it possible to estimate the amounts and ratios of key nutrients ingested (in GF parlance, this is known as an ‘intake target’). Third, and finally, chemical profiles of the animals can be conducted (e.g., body protein, sugar and lipid content can be measured; in the GF such measures are called ‘growth targets’). Linking intake targets and growth targets allows nutrient utilization to be assessed. The GF has proven powerful in manipulative research and has been extensively used in highly controlled laboratory conditions but only in very few observations of free-living animals in the wild (Raubenheimer 2011). The current study aims to apply, for the first time, the GF in field conditions, using a non-manipulative approach on a complex biological system that involves manipulation of the host-plant. This allows the understanding of the interplay between an insect and its host-plant under natural conditions, with a clear characterization of the amount and composition of food ingested by the insect, and the evaluation of related nutrient allocation strategies. However, such studies are challenging in endophagous insects as the system is difficult to manipulate due to (*i*) their peculiar lifestyle within plant tissues and (*ii*) the lack of artificial diet these organisms for which the microenvironment generated is also critical for their survival. For these reasons, the use of GF allows not only the understanding of nutrient regulation in a biological system for which no artificial diet is available but also to add more physiological realism to plant-insect nutritional ecology studies.

Due to the high nutritional constraints faced by permanent leaf-miner insects such as *P. blancardella*, we hypothesize that this insect has evolved multiple ways to deal with inadequate nutrition from the plant. This is expected to include pre- and post-ingestive mechanisms to control for leaf nutritional composition (Figure 1d), nutrient intake and/or allocation of ingested nutrients. Based on insect nutritional requirements and current knowledge on this biological system, we further hypothesize insect control of the leaf physiology to operate not only on sugars but also on proteins for the two larval feeding modes (Figure 1d). Collectively, our approach provides a field-based view of nutrient regulation in a complex tripartite biological system that involves manipulation of the host-plant. We end the discussion with a comparison of optimal protein-carbohydrate requirements for 117 insect species, which reinforces the hypothesis of a close association between *P. blancardella* and endosymbiotic bacteria for nutritional purposes.

## Material and methods

### Biological material

The experiments were conducted on *Malus domestica* (Borkh., 1803) (Rosaceae) apple-tree leaves naturally infected by the spotted tentiform leaf-miner, *Phyllonorycter blancardella* (Fabricius, 1781) (Lepidoptera: Gracillariidae). This endophytophagous insect, which lives concealed within plant tissues and feeds internally, is a polyvoltine leaf-mining microlepidopteran widely distributed in Europe. Adult females randomly lay eggs on the lower surface of green apple-tree leaves. When larvae hatch they directly enter into the leaf through the epidermis without contact with the ambient environment. Larvae must make the best of their mother’s choice, as they cannot abandon this leaf. There are some exceptions (Needham et al. 1928) – for example, the Diptera larva *Scaptomyza flava* (Whiteman et al. 2011), the Coleoptera larva *Neomycta rubida* (Martin 2010) and the micro-Lepidoptera *Coleophora klimeschiella* (Khan and Baloch 1976) – but generally this strategy is very rare, and restricted to certain groups.

Larval development for *P. blancardella* is divided into five instars, with two distinct feeding modes (see Body et al. 2015 for more details). During the three first developmental stages (L1-L2-L3), caterpillars define the outline of their mine by separating the two leaf integuments; this also acts to determine the total surface available to later stages. The process of separating the leaf integument generates fluids, on which the L1-L2-L3 stages feed; appropriately, these caterpillars are called fluid-feeding instars (in some of the literature they are called “sap-feeders” – Pottinger and LeRoux 1971) (Body et al. 2015). During the two last instars (L4-L5), larvae selectively consume mesophyll cells and are called tissue-feeders. The removal of mesophyll results in the formation of translucent patches commonly referred to as a “feeding window” (Pottinger and LeRoux 1971; Body et al. 2015).

Usually only one mine is found per leaf but higher population densities can lead up to three mines per leaf (Pottinger and LeRoux 1971). When two larvae accidentally join their mines, larval competition will lead to the development of only a single adult (Pottinger and LeRoux 1971). Mines are uniformly distributed in trees and can occur in every location on a leaf (Pottinger and LeRoux 1971). The last generation of insects (October-November) occupies leaves that are undergoing senescence. These leaves can turn from green to yellow, and in the absence of endosymbionts senescing yellow leaves do not support caterpillar development (Kaiser et al. 2010; Body et al. 2013; Gutzwiller et al. 2015).

Both green and yellow mined leaves (only one mine per leaf), and their respective unmined green and yellow controls (an adjacent neighboring leaf), were simultaneously collected in autumn (October-November; the last generation of *P. blancardella*), on 16-18 years old apple-trees (“Elstar” varieties), in Thilouze, France (47° 14’ 35” North, 0° 34’ 43” East); all collections took place between 8:00 am and 10:00 am. The synchronization of sampling is crucial as levels of sugars, for example, greatly vary during the day and between different seasons. This required collecting green and yellow leaves mined by fluid- and tissue-feeding larvae and their respective unmined controls simultaneously to make sure that any observed physiological differences were due to the impact of the leaf-miner on the plant, and not to phenological changes in the trees.

Collected leaves and associated larvae (*green leaves:* N = 15 for fluid-feeding instars, N = 27 for tissue-feeding instars; *yellow leaves:* N = 15 for fluid-feeding instars, N = 26 for tissue-feeding instars) were immediately dissected on ice. To study the spatial (mined *vs.* unmined areas) and temporal (senescence) variation of protein and soluble sugar concentrations, mined tissues (zone M; excluding the leaf-miner, and its faeces), ipsilateral tissues (zone UM^1^; leaf tissues on the same side of the main vein as the mine), and contralateral tissues (zone UM^2^; leaf tissues on the opposite side of the main vein as the mine) were dissected (Giron et al. 2007). Non-infected green and yellow leaves (zone UM^3^) were also collected to check whether leaf-miners can impact adjacent neighboring leaves through systemic effects (Giron et al. 2007; Body 2013; Body et al. 2013; Body et al. in prep). Dissected samples were then stored at −80*°*C until analysis.

### Nutrient quantification in leaf tissues

#### Sample preparation

Following lyophilization (primary desiccation of 1 hour at −10°C and 25 mbar, followed by a secondary desiccation at −76°C and 0.001 mbar overnight; Bioblock Scientific Alpha1-4LDplus lyophilizer), leaf samples were ground, in liquid nitrogen, to an extra-fine powder. Similar amounts of mined, ipsilateral, contralateral and non-infected plant tissues were used to allow qualitative and quantitative comparisons (Sartorius micro-balance model 1801-001, Sartorius SA, Palaiseau, France).

#### Nutrient extraction

Prior to colorimetric quantifications, chlorophyll and other pigments were removed from leaf tissues (5 mg) with acetone (100 %) until complete elimination of natural coloration. Soluble sugars and proteins were extracted with vortex agitation for 30 sec at room temperature in 1 mL aqueous methanol (80 %) (Fisher Scientific; Hampton, New Hampshire, USA). After centrifugation at 1500 rpm, soluble sugars and proteins in leaf tissues were colorimetrically measured following established protocols based on Anthrone (sugar) and Bradford’s (protein) reagents (Giron et al. 2002; Giron et al. 2007; Body 2013; Body et al. 2013; Body et al. in prep).

#### Total soluble sugar quantification

For each sample, 100 µL of initial aqueous methanol supernatant were transferred into a borosilicate tube (16 x 100 mm; Fisher Scientific; Hampton, New Hampshire, USA) and placed in a water bath at 90 °C to evaporate the solvent down to a few microlitres. After adding 1 mL of anthrone reagent, the tubes were placed in a water bath at 90 °C for 15 min, cooled down at 0 °C for 5 min, vortexed and then read in a spectrophotometer at 630 nm (DU®-64 spectrophotometer; Beckman, Villepinte, France). The anthrone reagent consisted of 1.0 g of anthrone (Sigma Aldrich; St. Louis, Missouri, USA) dissolved in 500 mL of concentrated sulfuric acid (Fisher Scientific; Hampton, New Hampshire, USA) added to 200 mL of MilliQ water (Merck Millipore; Billerica, Massachusetts, USA).

#### Total soluble protein quantification

For each sample, 150 µL of initial aqueous methanol supernatant were transferred into a hemolysis tube (10 x 75 mm; Fisher Scientific; Hampton, New Hampshire, USA), and 650 µL of saline solution 0.15 M (2.19 g sodium chloride in 250 mL of MilliQ water) were added onto each tube along with 200 µL of concentrated Bradford reagent (Bio-Rad, Hercules, California, USA). Samples were then vortexed and read in a spectrophotometer at 595 nm (DU®-64 spectrophotometer; Beckman, Villepinte, France).

#### Calibration curves

Calibration curves that allowed us to transform absorbance into concentrations were made with bovine serum albumin (BSA; Sigma-Aldrich, St. Louis, Missouri, USA) for proteins. For total soluble sugars, calibration curves were corrected for the underestimation of sugar alcohols using a sugar mixture (sorbitol, trehalose, sucrose, glucose and fructose; Sigma Aldrich; St. Louis, Missouri, USA) (Body 2013; Body et al. 2013) close to the composition of mined and unmined tissues both on green and on yellow leaves.

### Nutrient quantification in leaf tissues

Larvae collected from green and yellow leaf samples were weighed (Mettler-Toledo micro-balance model ME30, Mettler-Toledo, Viroflay, France); fluid-feeding instars, because of their very small size, were pooled while keeping separated larvae from green and yellow leaves (*green leaves:* N = 15; *yellow leaves:* N = 15). Proteins, soluble sugars, and lipids (Vanillin reagent) in larvae were colorimetrically quantified following Foray et al.’s protocol (Foray et al. 2012).

### Geometrical framework

The geometrical framework (GF) is a state-space modeling approach that explores how an animal solves the problem of balancing multiple nutritional needs in a multidimensional and variable environment (Raubenheimer and Simpson 1999). It treats an animal as living within a multidimensional nutrient space where there are as many axes as there are functionally relevant (fitness-affecting) nutrients. There are more than thirty required nutrients for most animals, but protein and carbohydrates are among the most important for herbivores because their concentrations in plants are highly variable and often limiting (Behmer and Joern 2008). This approach, developed under highly controlled lab conditions using artificial diets, provides a theoretical unifying framework leading to a deep understanding of behavioral and physiological mechanisms underlying nutritional homeostasis and how the nutritional environment of animals impact their performances (Raubenheimer et al. 2009; Simpson and Raubenheimer 2012).

The composition of leaf tissues consumed (protein and sugar amounts) determines the “nutritional landscape” used by insects, and is expressed as a percentage of the leaf dry weight (as average ± S.E.M.). The withdrawal of sugar-rich mesophyll tissues by leaf-mining insects, and the over-representation of sugar-free epidermis in the mined tissue samples, must be taken into account when comparing mined and unmined tissues. Thus, gravimetry was used to estimate the amount of mesophyll eaten by larvae, which in turn allowed us to correct biochemical data accordingly (see Supplement 1).

The amount of leaf tissues ingested (quantified by gravimetry; see Supplement 1), and the specific nutrient composition of these tissues, were used to estimate the amounts of protein and sugar ingested by leaf-mining larvae which correspond to their “intake target”. All data are expressed in µg (average ± S.E.M.).

The amounts of protein, sugar and lipid in caterpillars (body composition) were quantified to estimate their “growth target” and are expressed in µg (average ± S.E.M.). An exact determination of intake targets would require challenging leaf-mining larvae with various artificial diets which are not available for most of endophagous insects, including *P. blancardella*. However, in our attempt to use the GF in the field and on a biological system for which no artificial diet is available, we used the amount of leaf tissues ingested and the specific nutrient composition of these tissues to estimate the amounts of protein and sugar ingested by leaf-mining larvae.

Comparison between mined and unmined leaf tissues demonstrates pre-ingestive regulation of nutrient composition through manipulation of the host-plant macronutrient profile. In contrast, post-ingestive regulation of nutrients is shown through comparison of nutrient amounts ingested by the caterpillar (“intake target”) with the chemical composition of its body (“growth target”).

### Statistical analysis

Statistical analyses were performed using R version 3.2.1 and RStudio version 0.99.467 (The R Foundation for Statistical Computing, Vienna, Austria). When necessary, data were transformed using a Log10 transformation allowing the use of parametric statistical tests (Zar 2007). We used multivariate analyses of variance (MANOVA) to analyze (*i*) protein-sugar in plants, (*ii*) protein-sugar eaten by caterpillars, and (*iii*) protein-sugar and protein-lipid, in caterpillars. For all MANOVA analyses, we used the Pillai’s test statistic, which is considered to be the most robust to violations of assumptions (Scheiner 1993; Behmer and Joern 2008). Where significant effects were observed, post-hoc comparisons were performed.

Preliminary statistical analysis showed that there were no significant differences in the amounts of protein and sugar between unmined tissues (ipsilateral (UM^1^), contralateral (UM^2^) tissues) and non-infected leaf tissues from an adjacent neighboring leaf (UM^3^)) (MANOVA: *on green leaves*, F_2,124_ = 0.073, P = 0.928; *on yellow leaves*, F_2,121_ = 0.377, P = 0.686). This indicates that the leaf-miner effects on protein and sugar contents are restricted to the mine and do not impact adjacent leaves systemically, allowing us for the use ipsilateral (UM^1^) and contralateral (UM^2^) tissues as an unmined control. Additionally, the amounts of protein and sugar of unmined zones (UM^1^ and UM^2^) were identical (MANOVA: *on green leaves*, F_2,82_ = 0.026, P = 0.974; *on yellow leaves*, F_2,80_ = 0.162, P = 0.850). This allows the statistical comparison of mined (M) *vs.* unmined (UM) tissues (UM^1^ + UM^2^) in the result section.

## Results

### Leaf tissue soluble sugar and protein content: insect nutritional landscape

Here we made six comparisons. First, we contrasted the protein and soluble sugar profiles of unmined tissues from green (photosynthetically active tissue) and yellow leaves (senescing tissue). The protein and soluble sugar profiles of these two leaf types differed (Table 1a); green leaves, compared to yellow leaves, had significantly higher soluble sugar levels, but reduced protein levels (Figures 2a and 2b). Next, leaf protein and sugar amounts of mined tissues on green and yellow leaves was compared across a number of different conditions (Table 1b-f). The first comparison from these analyses is between mined tissues on green and yellow leaves. The protein-sugar profiles were significantly different (Table 1b), but this difference (for both feeding stages) was a function of soluble protein content, which was higher for mined tissues on yellow leaves (Figures 2a and 2b); soluble sugar content was similar in the two mined tissues of green and yellow leaves. The next two comparisons were between unmined and mined tissues, on both green and yellow leaves. On green leaves, a difference in the two tissue types was only observed during the fluid-feeding stage, with sugar levels being significantly reduced in mined tissues compared to unmined tissues (Table 1c; Figure 2). On yellow leaves, the nutrient profiles of mined and unmined tissues differed for both feeding stages (Table 1d). Here, sugar levels were significantly increased in mined tissues, but no differences in protein content were observed (Figures 2a and 2b). Two final comparisons were made. First, we compared the protein-sugar plant profile available to fluid and tissue-feeders on green leaves (Table 1e). Next, we compared protein-sugar plant profiles for fluid and tissue-feeders on yellow leaves (Table 1f). In both instances, the protein-sugar profiles of the mined tissues were similar for fluid and tissue-feeders.

**Figure 2.**
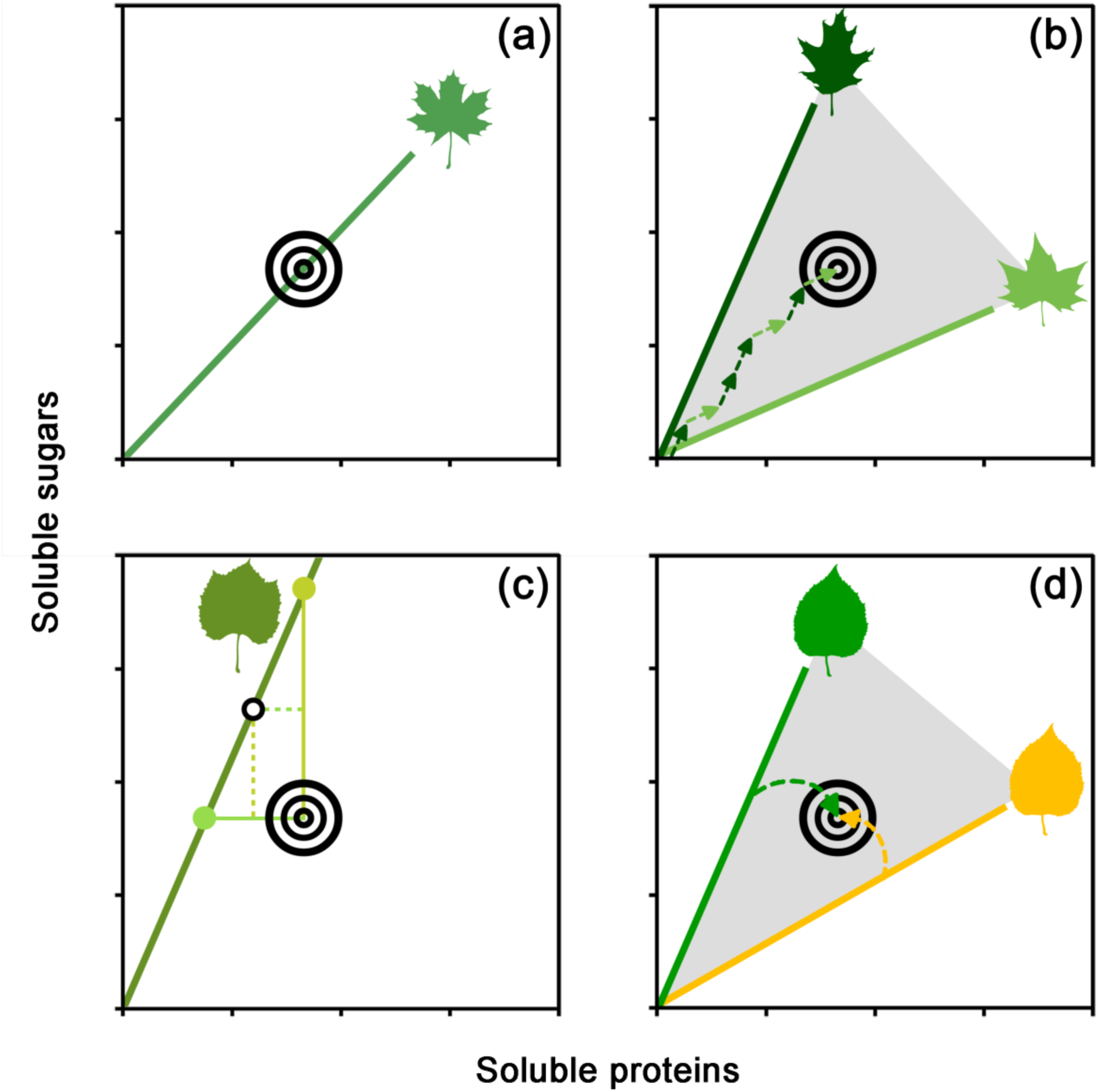
Nutrient landscapes for (a) early instar (fluid-feeders) and (b) late instar (tissue-feeders) caterpillars. For these two panels, the protein and soluble sugar content (expressed as the % dry mass of the tissue) of leaf tissues is shown. Square symbols represent unmined leaf tissues; triangles represent mined tissues for fluid-feeding caterpillars; circles represent mined tissues for tissue-feeding caterpillars. Closed symbols represent data obtained on green photosynthetically active leaves; open symbols represent data obtained on yellow senescing leaves which are no longer photosynthetically active. All data are presented as averages (± S.E.M). The shaded area comprised between the two extreme nutritional compositions of mined tissues depicts the most extensive nutrient landscape that *P. blancardella* larvae could encounter.

**Table 1.**
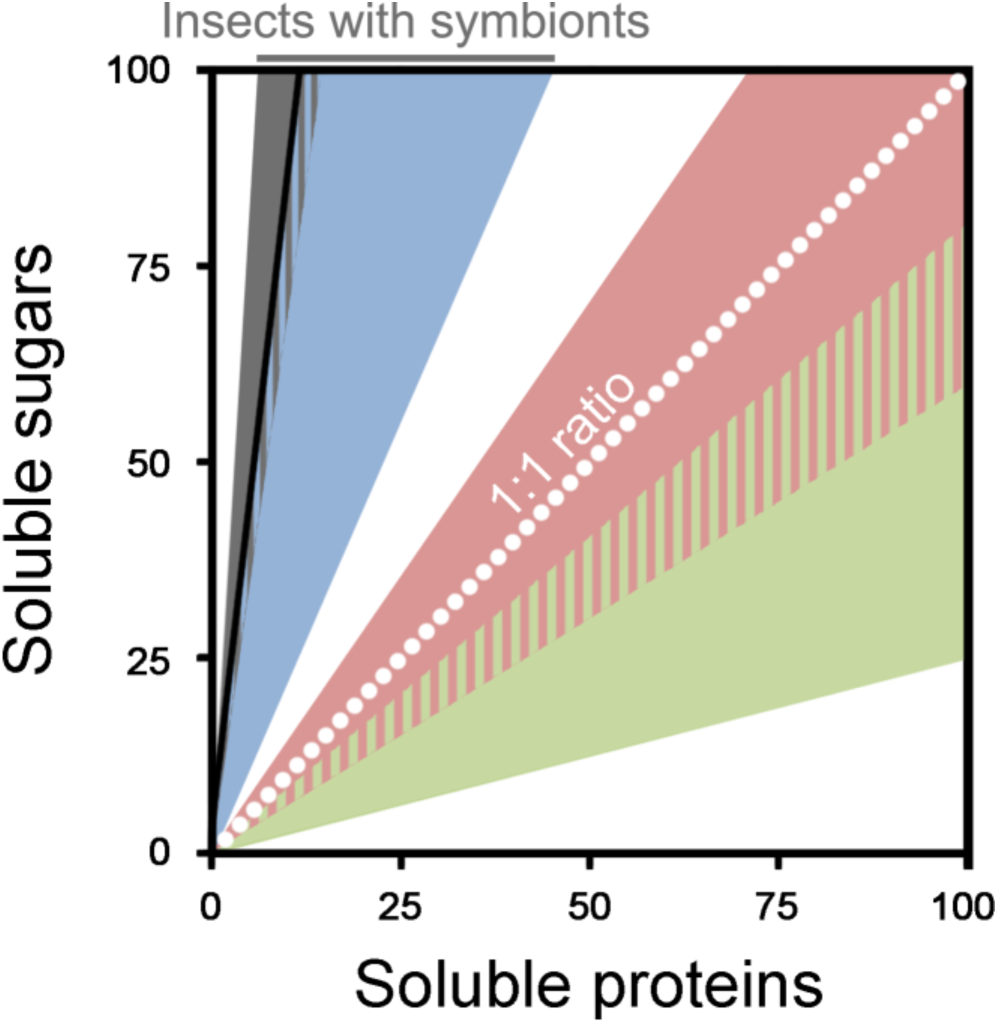
Statistical results from various comparisons related to plant nutritional content, plus caterpillar intake and growth targets. In all cases, MANOVA were conducted; for each comparison, univariate tests (shown under each comparison) were also conducted (ns = no significant difference, P > 0.05; * = P < 0.05; ** = P < 0.01; *** = P < 0.001). Caterpillar intake and growth targets were only measured for the tissue-feeding stage.

### Caterpillar intake and growth targets

The intake target, defined as the amount of sugars and proteins eaten, differed between green and yellow leaf tissue-feeding caterpillars (Table 1g); this was mostly a function of caterpillars on yellow leaves ingesting slightly more protein compared to caterpillars from green leaves (Table 1g; Figure 3a).

**Figure 3.**
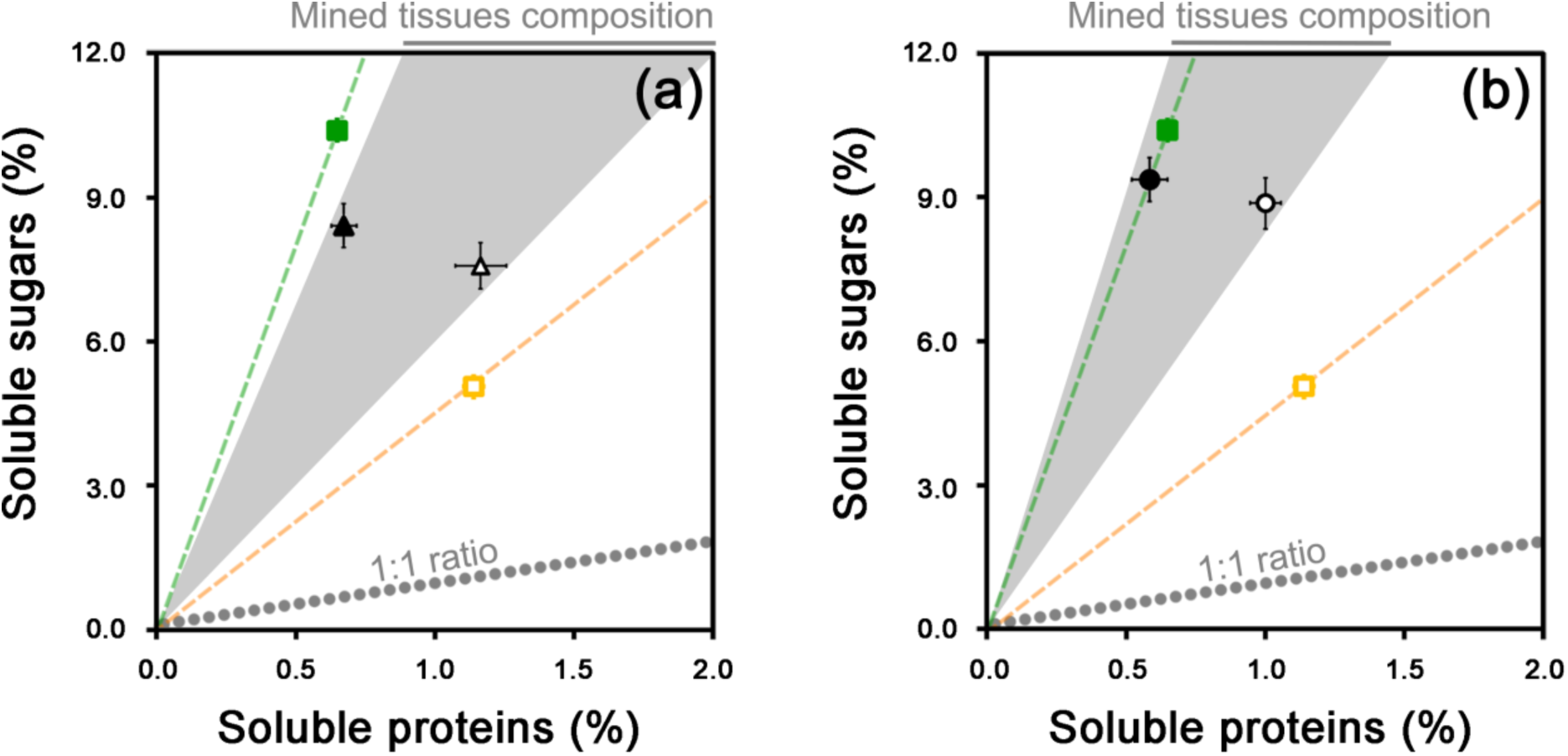
Panel (a) shows the soluble protein-sugar intake target (circles) for leaf-mining caterpillars (tissue-feeders) feeding on green and yellow leaves (expressed as µg per larva). The amount of leaf tissues ingested and the specific nutrient composition of these tissues were used to estimate the amounts of protein and sugar ingested by leaf-mining larvae which correspond to their “intake target”. Protein and soluble sugar contents of larvae on these tissues, which corresponds to their “growth targets”, are also shown (diamonds). Panel (b) shows protein and lipid body contents of caterpillars (diamonds) feeding on green and yellow leaves. Closed symbols represent data obtained for larvae feeding on green photosynthetically active leaves; open symbols represent data obtained for larvae feeding on yellow senescing leaves which are no longer photosynthetically active. All data are presented as averages (± S.E.M).

Two separate growth targets were analyzed. We found no difference in the protein and sugar amounts in caterpillars collected from green or yellow leaves (Table 1h; Figure 3a). Likewise, we found no difference in the protein and lipid amounts in caterpillars collected from green or yellow leaves (Table 1i; Figure 3b). Finally, body mass did not differ between tissue-feeding larvae feeding on green or yellow leaves (larvae on green leaves: 1.82 ± 0.16 mg; on yellow leaves: 1.40 ± 0.23 mg; Wilcoxon test: P = 0.161).

## Discussion

Biochemical analyses revealed an alteration of the nutritional composition of mined tissues when compared to uninfected control tissues (Figure 2). This effect was most notable on yellow leaves, where *P. blancardella* caterpillars maintained leaf sugar production in mined tissues (green-island); in unmined yellow tissues, sugar levels dropped. While protein is often considered the most limiting nutrient for insect herbivores, energy limitations can also constrain their performance (Behmer 2009, Roeder and Behmer 2014). Additionally, for most leaf-miner species, larvae live within the leaf throughout their development and are not able to switch between plants or leaves even under adverse autumnal conditions (Needham et al. 1928; Hering 1951; Schoonhoven et al. 2005; Body et al. 2015). Several leaf-miner species have been shown to prevent mined tissues from senescing through manipulation of the cytokinin profile of their host-plant (Engelbrecht et al. 1969; Giron et al. 2007; Kaiser et al. 2010; Body et al. 2013; Zhang et al. 2016, 2017). For *P. blancardella*, release of cytokinins to the plant is mediated by insect endosymbiotic bacteria (Kaiser et al. 2010; Body et al. 2013; Giron et al. 2013; Gutzwiller et al. 2015). Due to the positive impact of these phytohormones on plant sugar metabolism, leaf-miners can thus keep energy levels high (Body et al. 2013; Giron et al. 2013; Zhang et al. 2016; Body et al. in prep). For caterpillars feeding on senescing leaves, such manipulations create an enhanced nutritional micro-environment (Figure 2) (Body et al. 2013; Body et al. in prep). In fact, the nutritional landscape experienced by *P. blancardella* caterpillars feeding on senescing leaves is not dissimilar compared to caterpillars feeding on green leaves (Figure 2) (Raubenheimer and Simpson 1999; Behmer and Joern 2012; Body et al. 2013; Body et al. in prep). The fitness consequences of this manipulation are significant, because it allows the caterpillars to survive under adverse conditions, and to complete an additional generation (Kaiser et al. 2010; Body et al. 2013; Body et al. in prep). Interestingly, the ability of *P. blancardella* to prevent mined tissues from senescing, while keeping the leaf’s nutritional composition close to uninfected green leaves, is closely associated with the larval instars. Indeed, control of “nutritional homeostasis” of mined tissues is higher for late instars, which differ from younger larval instars in their feeding mode (fluid-*vs.* tissue-feeder; Figure 2a *vs.* 2b) (Body et al. in prep). Finally, our data also demonstrate that *P. blancardella* larvae are manipulating the nutritional composition of the green leaves. Here, though, caterpillars were only slightly suppressing soluble sugar levels. This generated a diet richer in relative protein content, compared to uninfected green control tissues. Two benefits are likely derived from decreasing the protein:sugar ratio: (*i*) faster development (Mattson 1980; Han et al. 2014; Larbat et al. 2016; Coqueret et al. 2017), and (*ii*) reduced costs associated with processing excess amounts of sugars (Warbrick-Smith et al. 2006; Coqueret et al. 2017).

Although the specific nutritional requirements of leaf-mining insects are unknown, under the assumption that nutritional regulatory mechanisms have been configured by natural selection to ensure that mined tissues provide the animal with the optimal amounts and balance of nutrients (Warbrick-Smith et al. 2006, 2009), the nutrient composition at the feeding site provides an indication of what macronutrient profile (nutrient landscape) is considered advantageous. But even within this nutrient landscape, the caterpillars still have some flexibility with respect to regulating, and utilizing their nutrient intake. Interestingly, and despite living in two different nutrient landscapes (Figure 2), caterpillars from green and yellow leaves showed very similar soluble protein:sugar intake targets (the position of the two intake targets differed, although no significant differences in protein or soluble sugar intake were detected; Figure 3a, Table 1g). In terms of the growth targets (body protein, sugar and lipid content; Figures 3a and 3b), no differences were observed for caterpillars feeding on the two different leaves. This final outcome is striking, considering that the leaf-miners develop on two very distinct nutritional environments. It is achieved through a three-step process: (*i*) caterpillars first use endosymbionts to initially modify the leaf physiology (a pre-ingestive mechanism) (Kaiser et al. 2010; Body et al. 2013), (*ii*) followed by self-selection to control nutrient intake (another pre-ingestive mechanism), (*iii*) and finally using post-ingestive mechanisms to differentially utilize ingested nutrients.

The larvae show high body protein content, despite the fact that the mined tissues they feed on tend to be low in protein (especially for caterpillars on green leaves). Some insects feeding on low-protein food sources employ symbiotic microorganisms that synthesize and provide key limiting amino acids, which can then be used as building blocks for animal generated protein (Chown and Nicolson 2004; Moran 2007; Douglas 2009, 2013; Gündüz and Douglas 2009; Frago et al. 2012). In *P. blancardella* larvae, symbiont-mediated post-ingestive mechanisms might allow for, or supplement, key amino acids involved in growth. Additionally, the low sugar, high lipid content measured in caterpillars indicates that *P. blancardella* larvae are converting sugars to lipids with high efficiency (Giron and Casas 2003; Behmer 2009).

Different animals utilize different sources of nutrients and evolve diverse life-history strategies, which suggest that intake targets move across evolutionary time-scales. A comparison of larval nutritional requirements of 117 species, from various insect groups (Figure 4; adapted from Simpson and Raubenheimer 1993; Behmer and Joern 2008), reveals that insects with the steepest target rail (lowest P:S ratio) are those with endosymbiotic bacteria that contribute to insect nitrogen metabolism. Based on our results, the intake target of *P. blancardella* leaf-mining larva nests within targets of insects with endosymbionts, rather than with other caterpillars (which to tend to have protein-biased intake targets; Behmer 2009). This reinforces the hypothesis of a close association between leaf-miners and endosymbiotic bacteria for nutritional purposes. A clear demonstration for a role of bacterial symbionts on the nutritional ecology of their leaf-mining insect-host would require global genomic approaches and/or obtaining a similar set of data from symbiont-free insects. However, non-infected *P. blancardella* do not occur naturally in the field and manipulative experiments under natural conditions allowing a simultaneous characterization of both plant and insect nutritional profiles have so far been unsuccessful. Nevertheless, *P. blancardella* relies on bacteria, most likely *Wolbachia*, to control the physiology of its host-plant through manipulation of phytohormone levels (Kaiser et al. 2010; Body et al. 2013). *Wolbachia* has also recently been shown to play a role as nutritional mutualist for bedbugs *Cimex lectularius* (Hosokawa et al. 2010) and could potentially have the ability to alter the gene expression of the plant in the *Diabrotica virgifera* / Maize system (Barr et al. 2010; but see Robert et al. 2013 for opposite results).

**Figure 4.**
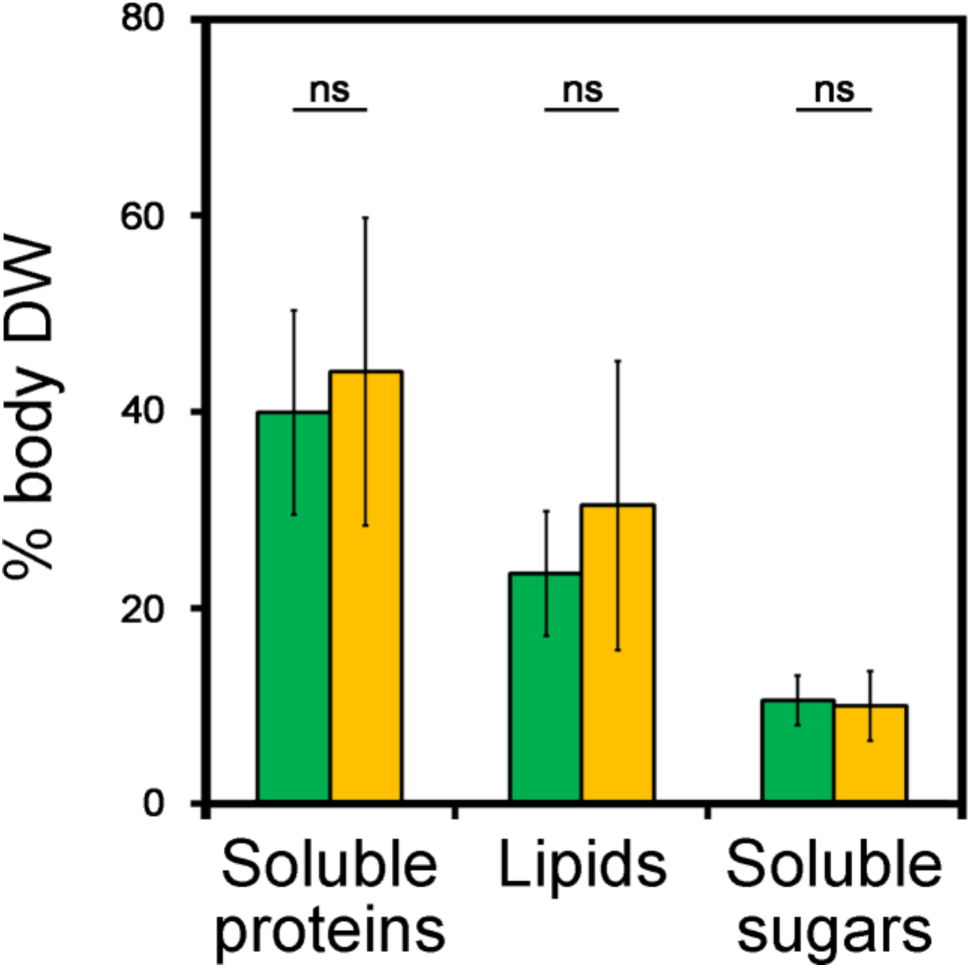
Intake target ranges for different insects (adapted from Simpson and Raubenheimer 1993; Behmer and Joern 2008). The shaded regions depict the range of intake targets for different insect groups: green = leaf-chewing caterpillars (N = 7), red = grasshoppers (N = 9), blue = insects (cockroaches and beetles) with symbionts (N = 5), black = aphids (N = 1), grey = *Phyllonorycter blancardella* (leaf-mining caterpillar from this current study).

Using the geometric framework for nutrition under natural field conditions allowed us to show that *P. blancardella* has multiple strategies to deal with a nutritionally variable and often suboptimal food supply in its host-plant. First, larvae manipulate the protein-sugar content of leaf tissues, and then use post-ingestive mechanisms to achieve similar body composition. Control of nutritional homeostasis of mined tissues is however stronger for late instars which differ from younger larval instars in their feeding mode. This advocate for a closer investigation of possible underlying mechanisms including variations of insect secretions, plant mechanical damages, and induced plant signalling responses over the course of the insect development. Manipulation of the leaf physiology also only impacts the leaf sugar profiles, most likely through a symbiont-mediated alteration of cytokinins. In contrast, insect control of protein metabolism only relies on post-ingestive mechanisms. Even though we have not been able to find a complete biosynthetic pathway for amino acids in *P. blancardella* bacterial symbiont (*Wolbachia*) (see Supplement 2), whether symbiotic microorganisms synthesize and provide key limiting amino acids in this system remains to be established. Using a similar approach on a larger range of organisms can extend the role of symbiotic microorganisms in insect nutrition beyond classical examples and holds considerable promise for promoting the study of field-based nutritional ecology.

## Acknowledgements

This study has been supported by the ANR project to DG ECOREN ANR-JC05-46491 and the Région Centre project to DG ENDOFEED 201000047141. We thank L. Ardouin for full access to his orchard, S. Venner for access to his lab, and W. Kaiser, E. Huguet, J.-P. Christidès, and the “Endofeed team” for helpful discussions.

