## Supplements for "Field application of the geometric framework reveals a multistep strategy of nutrient regulation in a leaf-miner"

**Supplement 1. Quantification of eaten tissues and correction for sugar content.**

The withdrawal of sugar-rich mesophyll tissues by leaf-mining insects, and the over-representation of sugar-free epidermis in the mined tissue samples, have been take into account when comparing mined and unmined tissues. Thus, gravimetry was used to estimate the amount of mesophyll eaten by larvae, which in turn allowed us to correct biochemical data accordingly.

Unmined areas similar to mined areas (Figure S1) were dissected in leaves for intermediate and late developmental stages (Figure S2). Following lyophilisation (Bioblock Scientific Alpha1-4LDplus lyophilizator), leaf samples were weighted (Mettler-Toledo micro-balance model ME30, Mettler-Toledo, Viroflay, France). Regression curves were used to correct biochemical data (Figure S3).

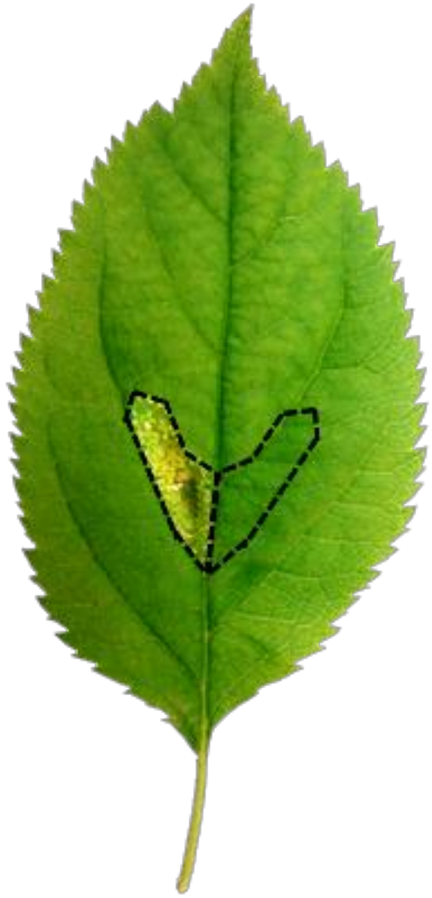

**Figure S1.** Similar areas of mined and unmined tissues dissected in apple-tree leaves.

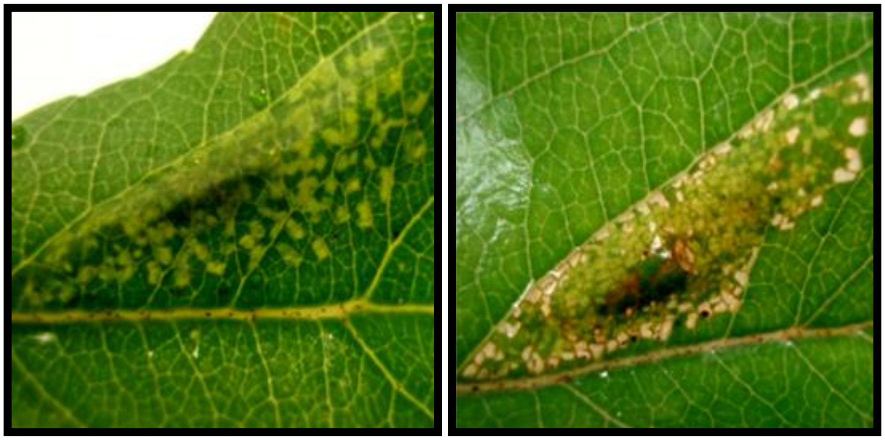

**Figure S2.** Mined tissues. Intermediate stage (left; L4 instar larvae) with spongy parenchyma and lower level palisade parenchyma cells consumed leading to light green patches. Late stage (right; L5 instar larvae) with spongy and palisade parenchyma cells consumed resulting in white and translucent patches where only the epidermis subsists.

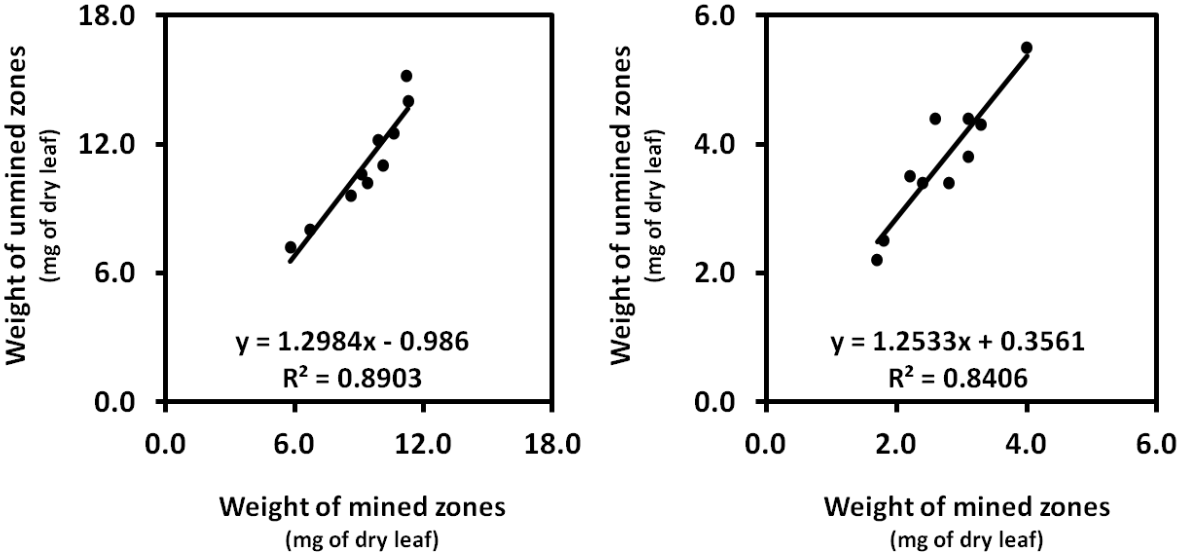


**Figure S3.** Regression curves used to correct data for intermediate (left) and late stage (right).

**Supplement 2. Amino acid biosynthesis pathways in *Phyllonorycter blancardella* endosymbiont, *Wolbachia pipientis*.**

The genome of *Wolbachia* associated to *P. blancardella* has not been sequenced yet and very few *Wolbachia* genomes are available. Therefore, a *Wolbachia* genome associated to *Drosophila melanogaster* (AE017196) was used as reference genome. This is likely that these two *Wolbachia* genomes from *Phyllonorycter blancardella* and *Drosophila melanogaster* share numerous pathways, even though a divergence for some functions cannot be excluded.

The comparison of established or predicted amino acid biosynthesis pathways in 8 well-known insect nutritional endosymbionts (Table S1) allowed to establish consensus gene sequences involved in amino acid synthesis. Among the 95 found genes, some of them coded for the same step but were specific of one insect symbiont.

| **Bacterial symbionts** | **Host-insects** |
|---|---|
| *Baumannia* | Sharpshooters |
| *Sulcia* | Sharpshooters |
| *Blattabacterium* | Cockroaches |
| *Blochmannia* | Ants |
| *Buchnera* | Aphids |
| *Carsonella* | Psyllids |
| *Ishikawaella* | Stinkbug |
| *Wigglesworthia* | Tsetse flies |
| ***Wolbachia*** | **Leaf-miners** |

**Table S1.** Host-insects and their associated bacterial symbionts used to establish consensus gene sequences involved in amino acid synthesis.

Based on already published genes known to be involved in amino acid biosynthesis (Table S2), protein sequences encoded by amino acid synthesis genes in symbiont genomes were identified using BLAST programs (Altschul et al., 1990) at National Centre for Biotechnology Information (NCBI) servers.

| Sequence | Name | Identifiant |
|---|---|---|
| NC_002528.1 | argA N-acetylglutamate synthase [Buchnera aphidicola str. APS (Acyrthosiphon pisum)] | ArgA [Buchnera aphidicola str. APS (Acyrthosiphon pisum)] |
| NC_002978.6 | argB acetylglutamate kinase [Wolbachia endosymbiont of Drosophila melanogaster] | ArgB [Wolbachia endosymbiont of Drosophila melanogaster] |
| NC_002528.1 | argC N-acetyl-gamma-glutamyl-phosphate reductase [Buchnera aphidicola str. APS (Acyrthosiphon pisum)] | ArgC [Buchnera aphidicola str. APS (Acyrthosiphon pisum)] |
| NC_002978.6 | argD acetylornithine transaminase protein [Wolbachia endosymbiont of Drosophila melanogaster] | ArgD [Wolbachia endosymbiont of Drosophila melanogaster] |
| NC_002528.1 | argE acetylornithine deacetylase [Buchnera aphidicola str. APS (Acyrthosiphon pisum)] | ArgE [Buchnera aphidicola str. APS (Acyrthosiphon pisum)] |
| NC_002528.1 | argF ornithine carbamoyltransferase subunit I [Buchnera aphidicola str. APS (Acyrthosiphon pisum)] | ArgF [Buchnera aphidicola str. APS (Acyrthosiphon pisum)] |
| NC_002528.1 | argG argininosuccinate synthase [Buchnera aphidicola str. APS (Acyrthosiphon pisum)] | ArgG [Buchnera aphidicola str. APS (Acyrthosiphon pisum)] |
| NC_002528.1 | argH argininosuccinate lyase [Buchnera aphidicola str. APS (Acyrthosiphon pisum)] | ArgH [Buchnera aphidicola str. APS (Acyrthosiphon pisum)] |
| NC_002528.1 | aroA 3-phosphoshikimate 1-carboxyvinyltransferase [Buchnera aphidicola str. APS (Acyrthosiphon pisum)] | AroA [Buchnera aphidicola str. APS (Acyrthosiphon pisum)] |
| NC_002528.1 | aroB 3-dehydroquinate synthase [Buchnera aphidicola str. APS (Acyrthosiphon pisum)] | AroB [Buchnera aphidicola str. APS (Acyrthosiphon pisum)] |
| NC_002528.1 | aroC chorismate synthase [Buchnera aphidicola str. APS (Acyrthosiphon pisum)] | AroC [Buchnera aphidicola str. APS (Acyrthosiphon pisum)] |
| NC_002528.1 | aroD 3-dehydroquinate dehydratase [Buchnera aphidicola str. APS (Acyrthosiphon pisum)] | AroD [Buchnera aphidicola str. APS (Acyrthosiphon pisum)] |
| NC_002528.1 | aroE shikimate 5-dehydrogenase [Buchnera aphidicola str. APS (Acyrthosiphon pisum)] | AroE [Buchnera aphidicola str. APS (Acyrthosiphon pisum)] |
| NC_002528.1 | aroH Trp-sensitive [Buchnera aphidicola str. APS (Acyrthosiphon pisum)] | AroH [Buchnera aphidicola str. APS (Acyrthosiphon pisum)] |
| NC_002528.1 | aroK shikimate kinase I [Buchnera aphidicola str. APS (Acyrthosiphon pisum)] | AroK [Buchnera aphidicola str. APS (Acyrthosiphon pisum)] |
| NC_002528.1 | asd aspartate-semialdehyde dehydrogenase [Buchnera aphidicola str. APS (Acyrthosiphon pisum)] | Asd [Buchnera aphidicola str. APS (Acyrthosiphon pisum)] |
| NC_010473.1 | asnA asparagine synthetase AsnA [Escherichia coli str. K-12 substr. DH10B] | AsnA [Escherichia coli str. K-12 substr. DH10B] |
| NC_004344.2 | asnB [Wigglesworthia glossinidia endosymbiont of Glossina brevipalpis] | AsnB [Wigglesworthia glossinidia endosymbiont of Glossina brevipalpis] |
| NC_002978.6 | aspC aspartate aminotransferase [Wolbachia endosymbiont of Drosophila melanogaster] | AspC [Wolbachia endosymbiont of Drosophila melanogaster] |
| NC_013364.1 | avtA valine--pyruvate transaminase [Escherichia coli O111:H- str. 11128] | AvtA [Escherichia coli O111-H- str. 11128] |
| NC_002528.1 | cysE serine acetyltransferase [Buchnera aphidicola str. APS (Acyrthosiphon pisum)] | CysE [Buchnera aphidicola str. APS (Acyrthosiphon pisum)] |
| NC_002528.1 | cysK cysteine synthase A [Buchnera aphidicola str. APS (Acyrthosiphon pisum)] | CysK [Buchnera aphidicola str. APS (Acyrthosiphon pisum)] |
| NC_002978.6 | dapA dihydrodipicolinate synthase [Wolbachia endosymbiont of Drosophila melanogaster] | DapA [Wolbachia endosymbiont of Drosophila melanogaster] |
| NC_002978.6 | dapB dihydrodipicolinate reductase [Wolbachia endosymbiont of Drosophila melanogaster] | DapB [Wolbachia endosymbiont of Drosophila melanogaster] |
| NC_006958.1 | dapC N-succinyldiaminopimelate aminotransferase [Corynebacterium glutamicum ATCC 13032] | DapC [Corynebacterium glutamicum ATCC 13032] |
| NC_002978.6 | dapD 2,3,4,5-tetrahydropyridine-2,6-carboxylate N-succinyltransferase [Wolbachia endosymbiont of Drosophila melanogaster] | DapD [Wolbachia endosymbiont of Drosophila melanogaster] |

| | | |
|---|---|---|
| NC_002978.6 | dapE succinyl-diaminopimelate desuccinylase [Wolbachia endosymbiont of Drosophila melanogaster] | DapE [Wolbachia endosymbiont of Drosophila melanogaster] |
| NC_002978.6 | dapF diaminopimelate epimerase [Wolbachia endosymbiont of Drosophila melanogaster] | DapF [Wolbachia endosymbiont of Drosophila melanogaster] |
| NC_013956.2 | gdhA Glutamate dehydrogenase [Pantoea ananatis LMG 20103] | GdhA [Pantoea ananatis LMG 20103] |
| NC_002978.6 | glnA glutamine synthetase [Wolbachia endosymbiont of Drosophila melanogaster] | GlnA [Wolbachia endosymbiont of Drosophila melanogaster] |
| NC_002978.6 | glyA serine hydroxymethyltransferase [Wolbachia endosymbiont of Drosophila melanogaster] | GlyA [Wolbachia endosymbiont of Drosophila melanogaster] |
| NC_002528.1 | hisA 1-(5-phosphoribosyl)-5-[(5-phosphoribosylamino)methylideneamino] imidazole-4-carboxamide isomerase [Buchnera aphidicola str. APS (Acyrthosiphon pisum)] | HisA [Buchnera aphidicola str. APS (Acyrthosiphon pisum)] |
| NC_002528.1 | hisB imidazole glycerol-phosphate dehydratase/histidinol phosphatase [Buchnera aphidicola str. APS (Acyrthosiphon pisum)] | HisB [Buchnera aphidicola str. APS (Acyrthosiphon pisum)] |
| NC_002528.1 | hisC histidinol-phosphate aminotransferase [Buchnera aphidicola str. APS (Acyrthosiphon pisum)] | HisC [Buchnera aphidicola str. APS (Acyrthosiphon pisum)] |
| NC_002528.1 | hisD histidinol dehydrogenase [Buchnera aphidicola str. APS (Acyrthosiphon pisum)] | HisD[Buchnera aphidicola str. APS (Acyrthosiphon pisum)] |
| NC_002528.1 | hisF imidazole glycerol phosphate synthase subunit HisF [Buchnera aphidicola str. APS (Acyrthosiphon pisum)] | HisF [Buchnera aphidicola str. APS (Acyrthosiphon pisum)] |
| NC_002528.1 | hisG ATP phosphoribosyltransferase [Buchnera aphidicola str. APS (Acyrthosiphon pisum)] | HisG [Buchnera aphidicola str. APS (Acyrthosiphon pisum)] |
| NC_002528.1 | hisH imidazole glycerol phosphate synthase subunit HisH [Buchnera aphidicola str. APS (Acyrthosiphon pisum)] | HisH [Buchnera aphidicola str. APS (Acyrthosiphon pisum)] |
| NC_006138.1 | hisHF bifunctional histidine biosynthesis protein (HisHF) [Desulfotalea psychrophila LSv54] | HisHF [Desulfotalea psychrophila LSv54] |
| NC_002528.1 | hisI bifunctional phosphoribosyl-AMP cyclohydrolase/phosphoribosyl-ATP pyrophosphatase protein [Buchnera aphidicola str. APS (Acyrthosiphon pisum)] | HisI [Buchnera aphidicola str. APS (Acyrthosiphon pisum)] |
| NC_002528.1 | ilvC ketol-acid reductoisomerase [Buchnera aphidicola str. APS (Acyrthosiphon pisum)] | IlvC [Buchnera aphidicola str. APS (Acyrthosiphon pisum)] |
| NC_002528.1 | ilvD dihydroxy-acid dehydratase [Buchnera aphidicola str. APS (Acyrthosiphon pisum)] | IlvD [Buchnera aphidicola str. APS (Acyrthosiphon pisum)] |
| NC_005061.1 | ilvE branched-chain amino acid aminotransferase [Candidatus Blochmannia floridanus] | IlvE [Candidatus Blochmannia floridanus] |
| NC_002528.1 | ilvH acetolactate synthase small subunit [Buchnera aphidicola str. APS (Acyrthosiphon pisum)] | IlvH [Buchnera aphidicola str. APS (Acyrthosiphon pisum)] |
| NC_002528.1 | ilvI acetolactate synthase large subunit [Buchnera aphidicola str. APS (Acyrthosiphon pisum)] | IlvI [Buchnera aphidicola str. APS (Acyrthosiphon pisum)] |
| NC_002253.1 | leuA 2-isopropylmalate synthase [Buchnera aphidicola str. APS (Acyrthosiphon pisum)] | LeuA [Buchnera aphidicola str. APS (Acyrthosiphon pisum)] |
| NC_002253.1 | leuB 3-isopropylmalate dehydrogenase [Buchnera aphidicola str. APS (Acyrthosiphon pisum)] | LeuB [Buchnera aphidicola str. APS (Acyrthosiphon pisum)] |
| NC_002253.1 | leuC isopropylmalate isomerase large subunit [Buchnera aphidicola str. APS (Acyrthosiphon pisum)] | LeuC [Buchnera aphidicola str. APS (Acyrthosiphon pisum)] |
| NC_002253.1 | leuD isopropylmalate isomerase small subunit [Buchnera aphidicola str. APS (Acyrthosiphon pisum)] | LeuD [Buchnera aphidicola str. APS (Acyrthosiphon pisum)] |
| NC_010471.1 | livA ABC-type branched-chain amino acid transport system, periplasmic component [Leuconostoc citreum KM20] | LivA [Leuconostoc citreum KM20] |
| NC_006576.1 | livE branched-chain amino acid aminotransferase [Synechococcus elongatus PCC 6301] | LivE [Synechococcus elongatus PCC 6301] |

| | | |
|---|---|---|
| NC_002528.1 | lysA diaminopimelate decarboxylase [Buchnera aphidicola str. APS (Acyrthosiphon pisum)] | LysA [Buchnera aphidicola str. APS (Acyrthosiphon pisum)] |
| NC_002528.1 | lysS lysyl-tRNA synthetase [Buchnera aphidicola str. APS (Acyrthosiphon pisum)] | LysS [Buchnera aphidicola str. APS (Acyrthosiphon pisum)] |
| NC_005061.1 | metB cystathionine gamma-synthase [Candidatus Blochmannia floridanus] | MetB [Candidatus Blochmannia floridanus] |
| NC_002978.6 | metC cystathionine beta-lyase [Wolbachia endosymbiont of Drosophila melanogaster] | MetC [Wolbachia endosymbiont of Drosophila melanogaster] |
| NC_002528.1 | metE 5-methyltetrahydropteroyltriglutamate--homocysteine S-methyltransferase [Buchnera aphidicola str. APS (Acyrthosiphon pisum)] | MetE [Buchnera aphidicola str. APS (Acyrthosiphon pisum)] .txt |
| NC_002528.1 | pheA chorismate mutase [Buchnera aphidicola str. APS (Acyrthosiphon pisum)] | PheA [Buchnera aphidicola str. APS (Acyrthosiphon pisum)] |
| NC_014125.1 | proA gamma-glutamyl phosphate reductase [Legionella pneumophila 2300/99 Alcoy] | ProA [Legionella pneumophila 2300-99 Alcoy] |
| NC_014125.1 | proB glutamate-5-kinase [Legionella pneumophila 2300/99 Alcoy] | ProB [Legionella pneumophila 2300-99 Alcoy] |
| NC_004344.2 | proC [Wigglesworthia glossinidia endosymbiont of Glossina brevipalpis] | ProC [Wigglesworthia glossinidia endosymbiont of Glossina brevipalpis] |
| NC_003062.2 | serA phosphoglycerate dehydrogenase [Agrobacterium tumefaciens str. C58] | SerA [Agrobacterium tumefaciens str. C58] |
| NC_002947.3 | serB phosphoserine phosphatase SerB [Pseudomonas putida KT2440] | SerB [Pseudomonas putida KT2440] |
| NC_002528.1 | serC phosphoserine aminotransferase [Buchnera aphidicola str. APS (Acyrthosiphon pisum)] | SerC [Buchnera aphidicola str. APS (Acyrthosiphon pisum)] |
| NC_002528.1 | thrA bifunctional aspartokinase I/homeserine dehydrogenase I [Buchnera aphidicola str. APS (Acyrthosiphon pisum)] | ThrA [Buchnera aphidicola str. APS (Acyrthosiphon pisum)] |
| NC_002528.1 | thrB homoserine kinase [Buchnera aphidicola str. APS (Acyrthosiphon pisum)] | ThrB [Buchnera aphidicola str. APS (Acyrthosiphon pisum)] |
| NC_002528.1 | thrC threonine synthase [Buchnera aphidicola str. APS (Acyrthosiphon pisum)] | ThrC [Buchnera aphidicola str. APS (Acyrthosiphon pisum)] |
| NC_002528.1 | trpA tryptophan synthase subunit alpha [Buchnera aphidicola str. APS (Acyrthosiphon pisum)] | TrpA [Buchnera aphidicola str. APS (Acyrthosiphon pisum)] |
| NC_012083.1 | trpAB tryptophane synthase [Thalassiosira pseudonana CCMP1335] | TrpAB [Thalassiosira pseudonana CCMP1335] |
| NC_002528.1 | trpB tryptophan synthase subunit beta [Buchnera aphidicola str. APS (Acyrthosiphon pisum)] | TrpB [Buchnera aphidicola str. APS (Acyrthosiphon pisum)] |
| NC_002528.1 | trpC bifunctional indole-3-glycerol phosphate synthase/phosphoribosylanthranilate isomerase [Buchnera aphidicola str. APS (Acyrthosiphon pisum)] | TrpC [Buchnera aphidicola str. APS (Acyrthosiphon pisum)] |
| NC_002528.1 | trpD anthranilate phosphoribosyltransferase [Buchnera aphidicola str. APS (Acyrthosiphon pisum)] | TrpD [Buchnera aphidicola str. APS (Acyrthosiphon pisum)] |
| NC_002252.1 | trpE anthranilate synthase component I [Buchnera aphidicola str. APS (Acyrthosiphon pisum)] | TrpE [Buchnera aphidicola str. APS (Acyrthosiphon pisum)] |
| NC_011989.1 | trpEG anthranilate synthase [Agrobacterium vitis S4] | TrpEG [Agrobacterium vitis S4] |
| NC_002252.1 | trpG2 anthranilate synthase component II [Buchnera aphidicola str. APS (Acyrthosiphon pisum)] | TrpG2 [Buchnera aphidicola str. APS (Acyrthosiphon pisum)] |
| NC_002252.1 | trpG anthranilate synthase component II [Buchnera aphidicola str. APS (Acyrthosiphon pisum)] | TrpG [Buchnera aphidicola str. APS (Acyrthosiphon pisum)] |
| NC_005061.1 | tyrA bifunctional chorismate mutase/prephenate dehydrogenase [Candidatus Blochmannia floridanus] | TyrA [Candidatus Blochmannia floridanus] |
| NC_000913.2 | tyrB tyrosine aminotransferase, tyrosine-repressible, PLP-dependent [Escherichia coli str. K-12 substr. MG1655] | TyrB [Escherichia coli str. K-12 substr. MG1655] |

**Table S2.** Sequences of genes coding for amino acid biosynthesis used to align against *Wolbachia* genome.

An exhaustive search of symbiont amino acid synthesis transcripts in the NCBI's databases was performed using the BLAST program (Altschul et al., 1990). The predicted translation of each full-length or partial sequence was checked and allowed the identification of top scoring matches for each nutritional symbiont. Alignment scores for each nutritional symbiont and each gene are presented in a Table 3 according to the following color key:

| Color key for alignment scores | | | | | |
|---|---|---|---|---|---|
| NO | <40 | 40-50 | 50-80 | 80-200 | >=200 |

| Gènes | *Baumannia* | *Blattabacterium* | *Blochmannia* | *Buchnera* | *Carsonella* | *Ishikawaella* | *Sulcia* | *Wigglesworthia* | *Wolbachia* |
|---|---|---|---|---|---|---|---|---|---|
| **AarAB** | | | | | | | | | |
| **AlaB** | | | | | | | | | |
| **ArgA** | | | black | red | | red | blue | | blue |
| **ArgB** | | green | | green | | blue | magenta | | red |
| **ArgC** | | magenta | black | red | | red | magenta | | black |
| **ArgD** | magenta | magenta | | red | | red | magenta | magenta | red |
| **ArgE** | | green | green | red | black | red | green | green | green |
| **ArgF** | green | magenta | red | red | magenta | red | magenta | magenta | green |
| **ArgG** | | magenta | black | red | red | red | magenta | | black |
| **ArgH** | blue | magenta | green | red | magenta | red | magenta | green | green |
| **ArgI** | green | green | red | red | green | red | green | green | green |
| **AroA** | red | green | red | red | green | red | green | red | black |
| **AroB** | red | green | red | red | green | red | green | red | blue |
| **AroC** | red | red | red | red | red | red | red | red | black |
| **AroD** | green | green | green | red | green | green | green | green | black |
| **AroE** | magenta | green | red | red | | red | black | magenta | |
| **AroF** | red | black | red | red | magenta | | red | red | |
| **AroG** | black | red | black | black | black | | red | black | black |
| **AroH** | red | magenta | red | red | magenta | red | | red | |
| **AroK** | red | magenta | red | red | black | red | magenta | red | black |
| **Asd** | black | magenta | red | red | red | red | magenta | red | magenta |
| **AsnA** | | black | | black | | | | | black |
| **AsnB** | | black | black | | | | | red | black |
| **AspB** | black | magenta | green | green | black | blue | magenta | black | magenta |
| **AspC** | blue | red | green | blue | | blue | red | black | red |
| **AstC** | magenta | magenta | | red | | red | magenta | magenta | red |

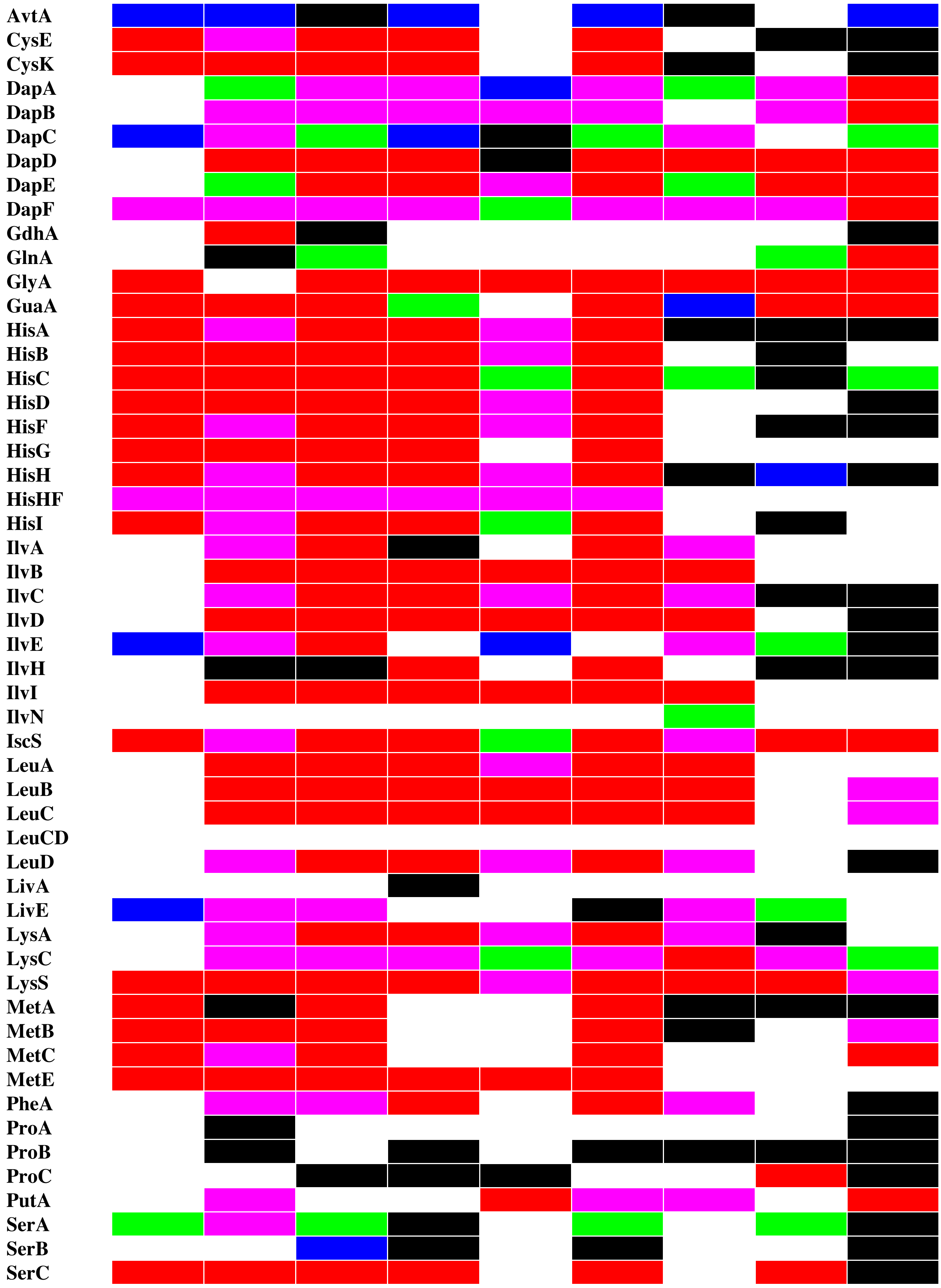
AvtA
CysE
CysK
DapA
DapB
DapC
DapD
DapE
DapF
GdhA
GlnA
GlyA
GuaA
HisA
HisB
HisC
HisD
HisF
HisG
HisH
HisHF
HisI
IlvA
IlvB
IlvC
IlvD
IlvE
IlvH
IlvI
IlvN
IscS
LeuA
LeuB
LeuC
LeuCD
LeuD
LivA
LivE
LysA
LysC
LysS
MetA
MetB
MetC
MetE
PheA
ProA
ProB
ProC
PutA
SerA
SerB
SerC

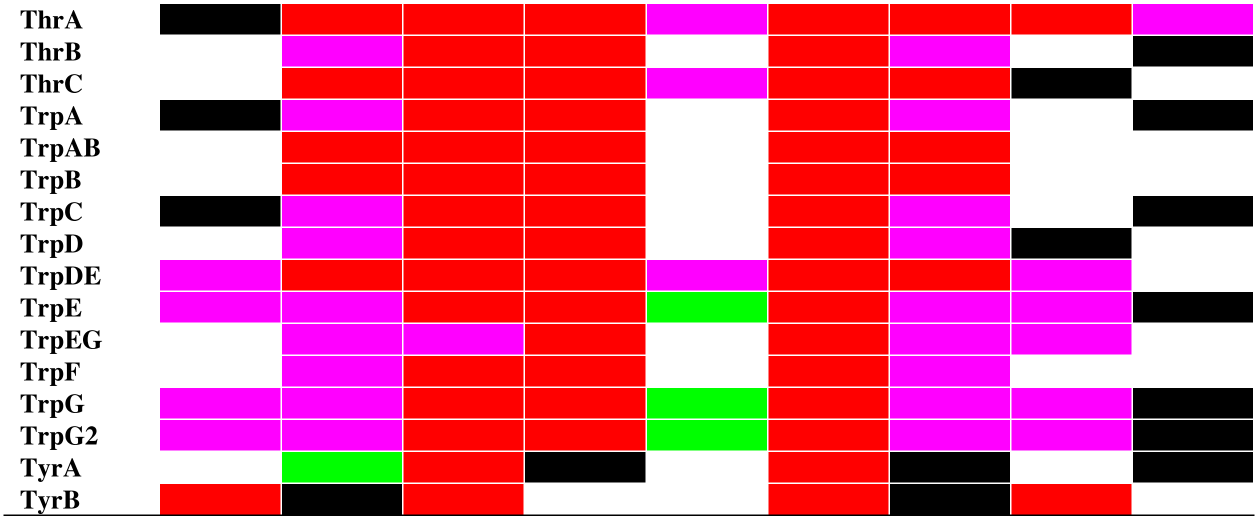


**Table S3.** Alignment scores for each bacterial symbiont and each gene coding for amino acid biosynthesis are summarized in this table.

An exhaustive search of symbiont amino acid synthesis transcripts in the NCBI's databases was performed using the BLAST program (Altschul et al., 1990). The predicted translation of each full-length or partial sequence was checked and allowed the identification of top scoring matches for each nutritional symbiont.

Genes coding for amino acid biosynthesis steps that where present in *Wolbachia* genome are summarized on the consensus gene sequences established from publications on amino acid biosynthetic pathways in insect-associated nutritional bacterial symbionts. Alignment scores for each gene are represented according to the following key:

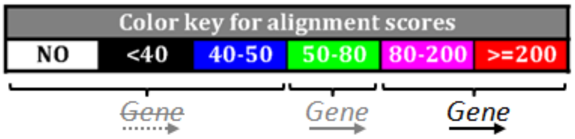

### a. Essential amino acid biosynthetic pathways

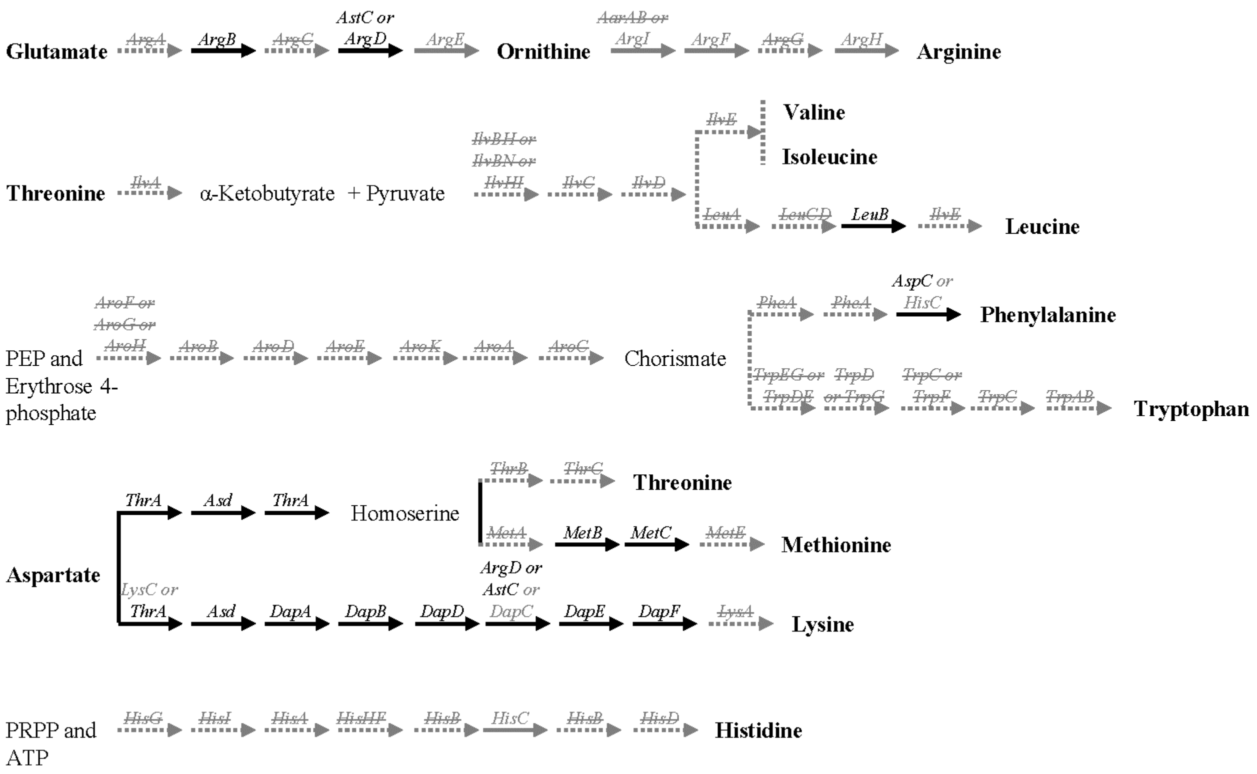


### b. Non-essential amino acid biosynthetic pathways

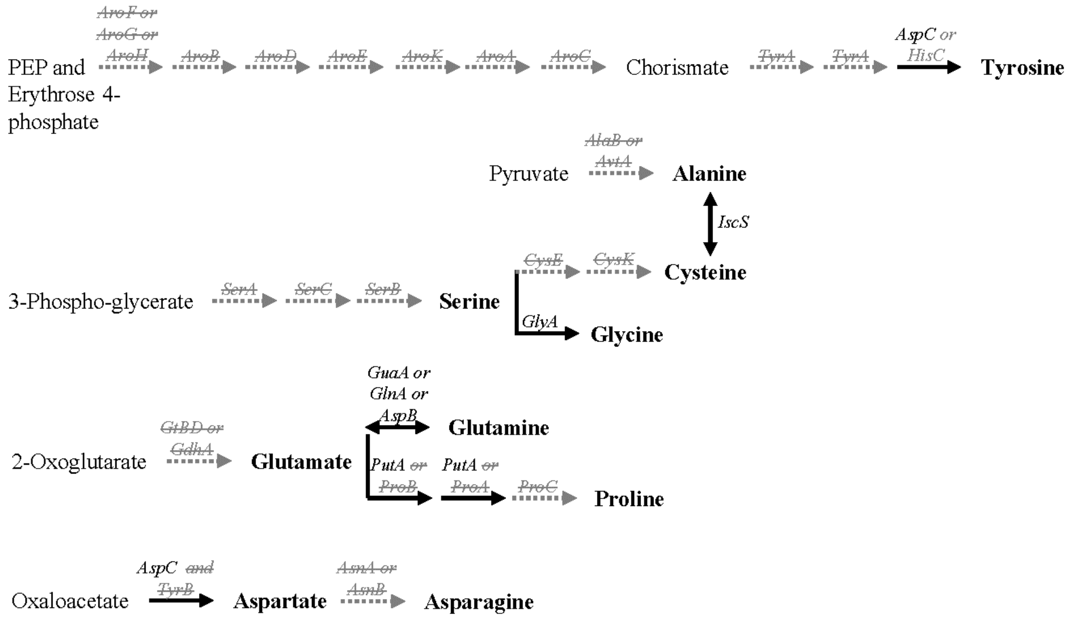
